# Integrated Omics Approach to Delineate the Mechanisms of Doxorubicin-Induced Cardiotoxicity

**DOI:** 10.1101/2025.06.24.661365

**Authors:** Mohamed S. Dabour, Ibrahim Y. Abdelgawad, Bushra Sadaf, Mary R. Daniel, Marianne K.O. Grant, Anne H. Blaes, Pamala A. Jacobson, Beshay N. Zordoky

## Abstract

Doxorubicin (DOX) is an effective chemotherapy whose clinical utility is limited by cardiotoxicity. To investigate underlying mechanisms, we employed a multi-omics approach integrating transcriptomic and proteomic profiling leveraging established mouse models of chronic DOX-induced cardiotoxicity. Five-week-old male mice received weekly DOX (4 mg/kg) or saline injections for six weeks, with heart tissues harvested 4 days post-treatment. Differentially expressed genes (DEGs) and proteins (DEPs) were identified by bulk RNA-seq and proteomics, validated via qPCR and western blot, respectively Key DEPs were validated in plasma samples from DOX-treated breast cancer patients. Additionally, a temporal comparison was conducted between DEPs in the mice hearts 4 days and 6 weeks post-DOX. RNA-seq revealed upregulation of stress-responsive genes (*Phlda3, Trp53inp1*) and circadian regulators (*Nr1d1*), with downregulation of *Apelin* and *Cd74*. Proteomics identified upregulation of serpina3n, thrombospondin-1, and epoxide hydrolase 1. Plasma SERPINA3 concentrations were significantly elevated in breast cancer patients 24 hours post-DOX. Gene set enrichment analysis (GSEA) revealed upregulated pathways including p53 signaling, apoptosis, and unfolded protein response. Integrated omics analysis revealed 2,089 gene-protein pairs. GSEA of concordant gene-protein pairs implicated p53 signaling, apoptosis, and epithelial-mesenchymal transition in upregulated pathways, while oxidative phosphorylation and metabolic pathways were downregulated. Temporal comparison with a delayed timepoint (6 weeks post-DOX) uncovered dynamic remodeling of cardiac signaling, with early response dominated by inflammatory and apoptotic responses, and delayed response marked by cell cycle and DNA repair pathway activation. This integrated-omics study reveals key molecular pathways and temporal changes in DOX-induced cardiotoxicity, identifying potential biomarkers for future cardioprotective strategies.

## 1. Introduction

Doxorubicin (DOX), an anthracycline antibiotic, stands as a cornerstone chemotherapeutic agent, demonstrating remarkable efficacy against a variety of adult and pediatric malignancies, including leukemias, lymphomas, sarcomas, and solid tumors such as breast cancer (1). Its potent anti-neoplastic activity stems primarily from its ability to intercalate into DNA, inhibit topoisomerase II, and generate reactive oxygen species (ROS), ultimately leading to cancer cell death (2). Despite its clinical utility, the clinical application of DOX is significantly constrained by dose-dependent, cumulative cardiotoxicity (1). This cardiotoxicity can manifest acutely during or shortly after treatment, or present later as chronic cardiomyopathy, potentially leading to irreversible heart failure years after therapy completion (1). The risk of developing cardiac dysfunction increases substantially with higher cumulative doses, posing a significant challenge for long-term cancer survivors (3).

The precise molecular mechanisms underlying DOX-induced cardiotoxicity are complex and not yet fully elucidated. While mitochondrial dysfunction, excessive ROS production, senescence, impaired calcium handling, and apoptosis have been implicated as key contributors (4, 5), a complete understanding of the intricate signaling pathways and cellular processes involved remains elusive. This knowledge gap highlights a critical need for comprehensive, unbiased investigations to unravel the molecular intricacies of DOX-induced cardiac dysfunction. Such understanding is imperative for developing effective strategies to predict, prevent, or mitigate this serious complication, thereby improving the safety profile of DOX and enhancing the quality of life for cancer patients.

Recent advancements in high-throughput ‘omics’ technologies, notably proteomics and transcriptomics, offer powerful, unbiased platforms for exploring complex biological processes like drug-induced organ toxicity (6). Proteomics allows for the large-scale identification and quantification of proteins within a specific tissue or biological system, providing insights into cellular functions, signaling pathways, and post-translational modifications altered by drug exposure (7). Similarly, transcriptomics, often performed via RNA sequencing (RNA-seq), enables the comprehensive profiling of gene expression changes, revealing alterations in transcriptional regulation and identifying genes and pathways affected by the drug (7). Integrating data from both proteomic and transcriptomic analyses can provide a more comprehensive and systems-level understanding of the molecular perturbations induced by DOX in the heart, potentially uncovering novel mechanisms and identifying key molecular players that might be missed by single-omic studies (8).

Therefore, the primary objective of the current study was to employ a multi-omics strategy, integrating cardiac proteomics and transcriptomics, to comprehensively delineate the molecular mechanisms underlying DOX-induced chronic cardiotoxicity. Using heart tissues from our previously published study of a chronic DOX-induced cardiotoxicity model in C57BL/6N mice (9), we aimed to identify key differentially expressed proteins (DEPs) and genes (DEGs), map the associated molecular pathways, and validate significant findings. Furthermore, we sought to assess the translational relevance of key identified biomarkers by examining their levels in plasma samples from breast cancer patients undergoing DOX-based chemotherapy. Additionally, we sought to characterize the temporal evolution of DOX-induced cardiotoxicity by comparing early (4 days post-DOX) and delayed (6 weeks post-DOX) proteomics data from our previously published study (10). By integrating these multi-level molecular data, we aimed to provide novel insights into the pathophysiology of DOX cardiotoxicity and identify potential therapeutic targets or biomarkers for early detection and intervention.

## 2. Methods

### 2.1. Animal Model and Doxorubicin Treatment

All animal procedures were approved by the Institutional Animal Care and Use Committee at the University of Minnesota (Protocol ID: 2106–39176 A) and conducted in accordance with relevant guidelines. The samples used in this study were derived from our previously published work (9, 10). Schematic design of the experiment is represented in Figure 1A. Five-week-old male C57BL/6N mice were allowed to acclimatize for one week prior to the experiment. Mice were then randomized into two groups: a control group receiving saline and a treatment group receiving DOX. DOX or saline was administered via intraperitoneal (IP) injection at a dose of 4 mg/kg body weight once weekly for a total duration of six weeks (n = 9 per group). Cardiac function was evaluated by echocardiography 3 days following the last injection of DOX and previously reported (9). Four days following the last DOX injection, mice were euthanized humanely by decapitation under isoflurane anesthesia, and organs were harvested. Organs were rinsed in ice-cold phosphate-buffered saline (PBS), flash-frozen in liquid nitrogen, and stored at – 80 ^◦^C until subsequent analysis.

**Figure 1.**
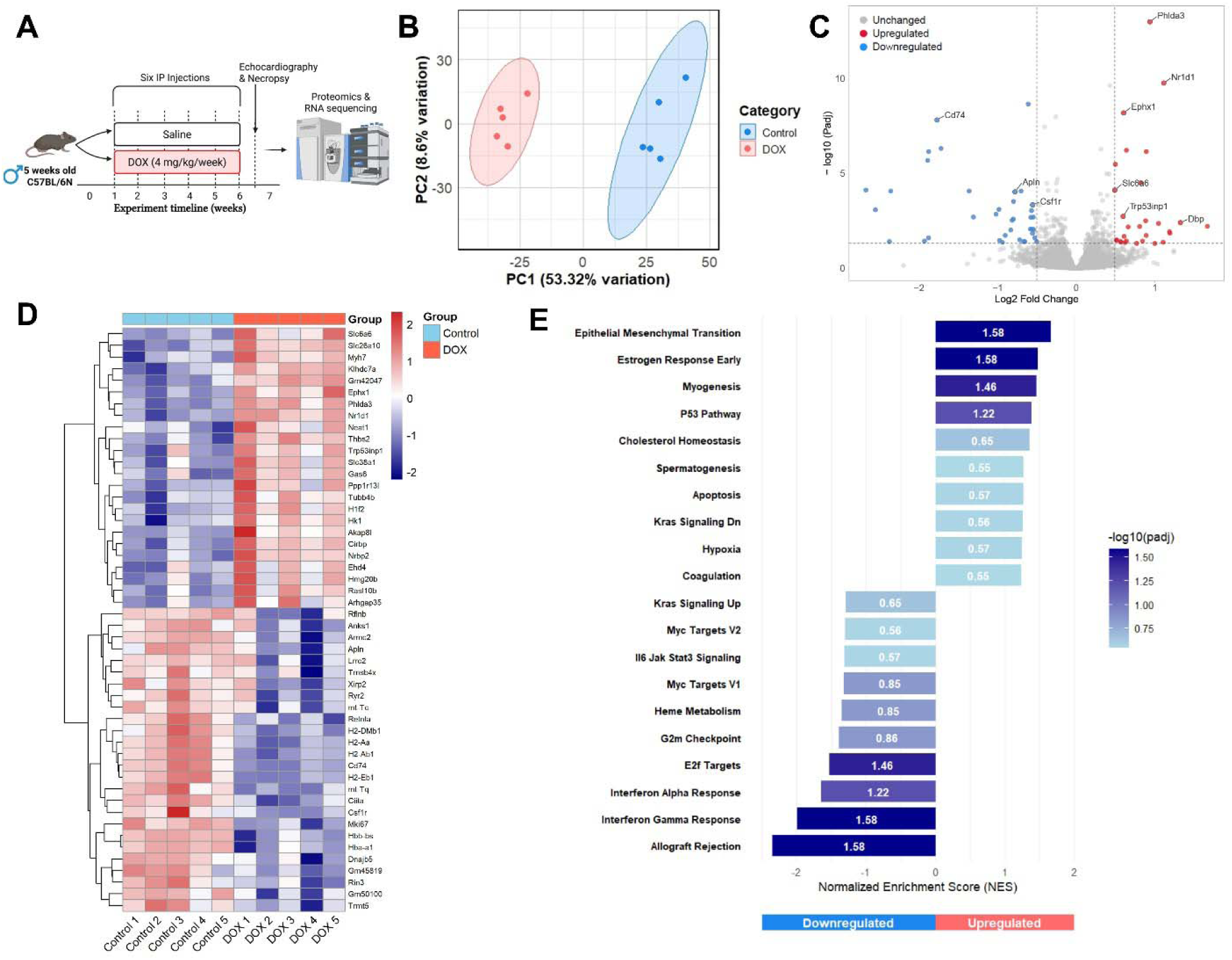
Experimental design and Transcriptomic Alterations Induced by DOX in Mouse Cardiac Tissue. (A) Experimental design for doxorubicin-induced cardiotoxicity model (B) Principal Component Analysis (PCA) plot showing distinct clustering of saline (Control, blue) and DOX-treated (red) samples. (C) Volcano plot showing differentially expressed genes (DEGs). Red dots indicate significantly upregulated genes, blue dots indicate significantly downregulated genes (based on |log_J fold change| > 0.5 and adjusted p-value < 0.05). Key DEGs are labeled. (D) Heatmap displaying relative expression levels (scaled) of the top 50 significantly DEGs across individual saline and DOX samples. Red indicates higher expression; blue indicates lower expression. (E) Gene Set Enrichment Analysis (GSEA) results showing selected significantly enriched pathways. Bars represent Normalized Enrichment Score (NES); positive NES indicates upregulation in DOX group, negative NES indicates downregulation. Color intensity corresponds to significance (-log10 adjusted p-value).

### 2.2. Bulk RNA-seq Analysis

RNA-seq analysis was performed on heart tissue samples (N=5 per group) with RNA-seq and initial bioinformatic processing were performed by Novogene Corporation (Sacramento, CA). Messenger RNA was purified from total RNA using poly-T oligo-attached magnetic beads, followed by fragmentation and cDNA synthesis using random hexamer primers. Two library types were constructed: non-directional libraries using dTTP and directional libraries using dUTP, both processed through end repair, A-tailing, adapter ligation, size selection, and amplification. Libraries were quantified using Qubit and real-time PCR before pooling and sequencing on Illumina platforms. The bioinformatics pipeline included quality control of raw reads using fastp software to remove adapter sequences and low-quality reads. Clean reads were aligned to the reference genome using Hisat2 v2.0.5, selected for its ability to generate splice junction databases from gene model annotations. Gene expression was quantified using featureCounts v1.5.0-p3 with FPKM normalization, and differential expression analysis was performed using DESeq2 (for samples with biological replicates) or edgeR (without replicates), with p-value adjustment using the Benjamini-Hochberg method and significance threshold of p≤0.05.

### 2.3. RNA extraction and Real-time PCR (qPCR)

Total RNA was isolated from cardiac tissue (20 mg) via homogenization in 300 μL of Trizol reagent (Life Technologies, Carlsbad, CA) in accordance with the manufacturer’s established protocol. RNA concentration and purity were assessed spectrophotometrically using a NanoDrop Lite instrument (Thermo Fisher Scientific, Waltham, MA) by measuring absorbance at 260 nm. Subsequently, first-strand cDNA synthesis was performed using 1.5 μg of purified total RNA with the Applied Biosystems high-capacity cDNA reverse transcription kit (Thermo Fisher Scientific) following the manufacturer’s specifications. Quantitative assessment of target gene expression was conducted via real-time PCR on a QuantStudio 5 platform utilizing SYBR Green chemistry (Thermo Fisher Scientific). Thermal cycling parameters consisted of initial denaturation at 95°C for 10 minutes, followed by 40 amplification cycles comprising denaturation at 95°C for 15 seconds and annealing/extension at 60°C for 1 minute. Oligonucleotide primer sequences employed for amplification are detailed in Table 1. Target gene expression was normalized to 18S ribosomal RNA as an endogenous reference and quantified relative to control samples using the comparative threshold cycle (ΔΔCT) methodology. Primer specificity and amplicon purity were verified through melting curve analysis of the terminal PCR products.

**Table 1.**
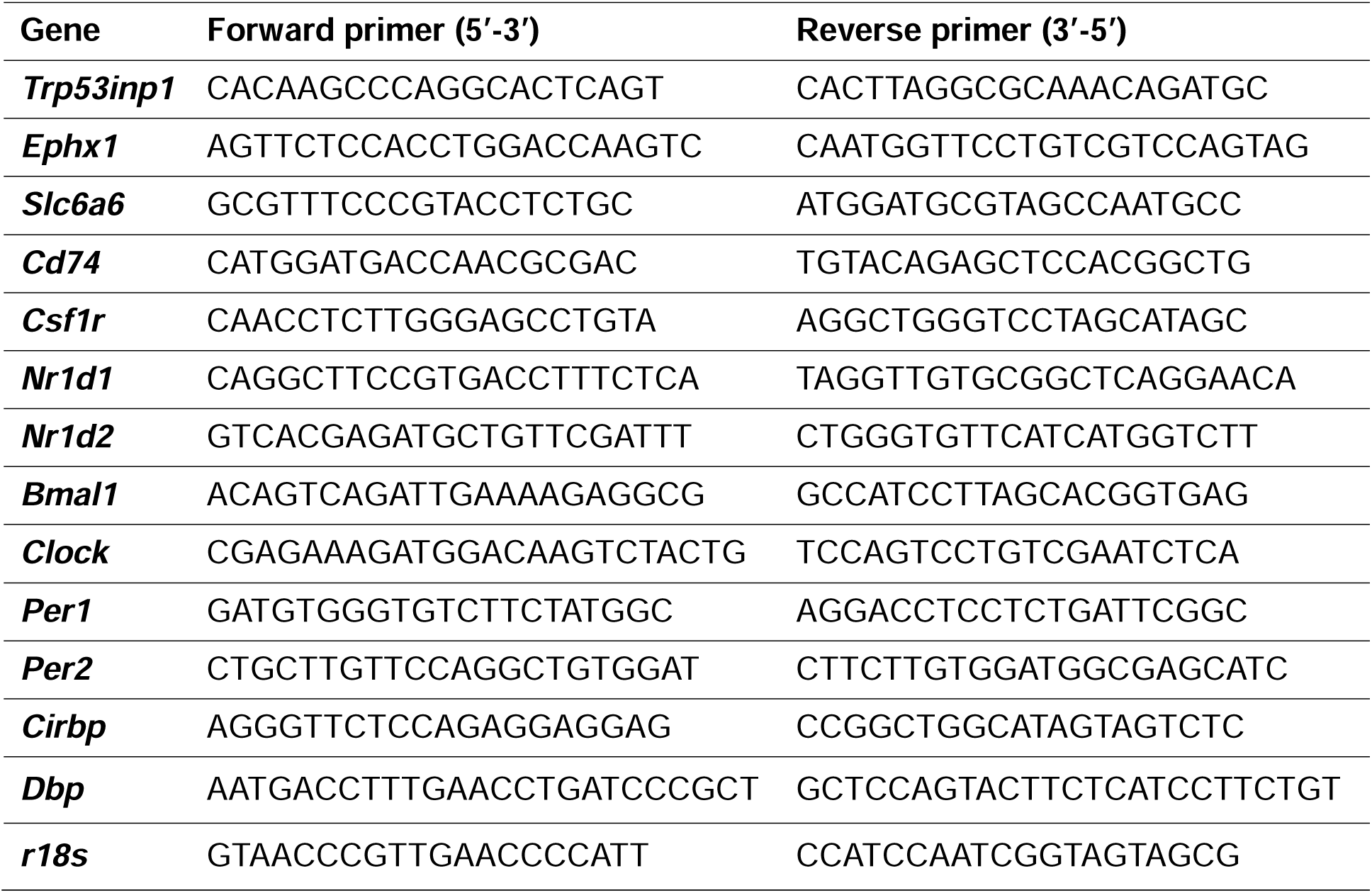
Primers used in this study.

### 2.4. Proteomics Sample Preparation and Analysis

Proteomics sample preparation and analysis (N=8 per group) was described previously (10). Following cardiac protein extraction, 25 μg protein aliquots were diluted fivefold with water before overnight trypsin digestion (1:40 ratio, 16 h, 37°C). Digested samples were acidified to 0.2% formic acid, purified using Waters Oasis MCX Prime cartridges, and resuspended in 0.1 M triethylammonium bicarbonate. For multiplexed quantification, 20 μg of each sample was labeled using TMTpro™ 16plex Isobaric Label Reagent according to manufacturer’s protocol (Thermo Fisher Scientific), combined, and purified using Oasis C18 solid phase extraction. Labeled peptides were fractionated by high pH C18 reversed-phase chromatography. Fractions were strategically concatenated into three groups (“early”, “middle”, and “late”) to create 10 final fractions, which were subsequently lyophilized and cleaned using the Stop and Go Extraction Protocol (STAGE tip). Raw data were analyzed using Proteome Discoverer software (version 2.2, Thermo Fisher Scientific). DEPs were identified using a threshold of false discovery rate (FDR) < 0.05. Statistical significance was determined using background t-tests with Benjamini-Hochberg multiple correction.

### 2.5. Protein Extraction and Western Blotting

Following euthanasia, cardiac tissues were rapidly snap-frozen in liquid nitrogen. Protein extraction from the frozen heart samples was performed using a previously established protocol (9). Total protein concentrations were quantified using the Pierce Bicinchoninic Acid (BCA) protein assay kit (Thermo Fisher Scientific). Equal amounts of protein (25 μg per sample) were separated by SDS–PAGE and transferred onto 0.2-μm nitrocellulose membranes. Immunoblotting was carried out using primary goat antibody against serpina3n (catalog # AF4709, R&D systems, Minneapolis, MN), and rabbit antibody against alpha-tubulin (catalog #2144, Cell Signaling, Danvers MA). Secondary antibodies included HRP-conjugated goat anti-rabbit and donkey anti-goat antibodies (catalog #111–035–144 and #705–035–147, respectively; Jackson ImmunoResearch, West Grove, PA). Protein band intensities were quantified using ImageJ software (NIH, Bethesda, MD), and α-tubulin was used as a loading control for normalization.

### 2.6. Bioinformatic Analyses

Bioinformatic analyses were performed to interpret the multi-omics data. Principal Component Analysis (PCA) was conducted on both transcriptomic and proteomic datasets using the FactoMineR R package after centering and scaling the data. Samples were visualized using ggplot2 with 95% confidence ellipses to indicate intra-group variability. Volcano plots were generated using VolcaNoseR with thresholds of |logLJ fold change| > 0.5 and adjusted p-value < 0.05. The top 50 DEGs and DEPs were selected based on adjusted p-values (FDR). Differential expression heatmaps for these top 50 DEGs and DEPs were generated using the pheatmap R package. Expression values were logLJ-transformed and standardized using row-wise z-scores prior to visualization. Gene set enrichment analysis (GSEA) was performed using the fgsea R package (v1.26.0) with gene sets from the Hallmark collection of the Molecular Signatures Database (MSigDB, version 2023.1). Only pathways containing 10-300 genes were included, and enrichment scores were computed using 1,000 permutations to calculate normalized enrichment scores (NES) and adjusted p-values. Top enriched pathways were visualized as bar plots annotated with –logLJLJ(padj) values.

For integrative analysis of transcriptomic and proteomic data, logLJ fold changes were extracted from both datasets and merged using gene symbols as identifiers. Genes were classified as concordant if the direction of fold change matched across platforms. A bubble plot was generated to visualize transcript-protein relationships, with point size representing supporting evidence and color indicating concordance. A subset of top genes was annotated using ggrepel, and Pearson correlation was calculated to evaluate the transcript-protein relationship. To compare early (4-day) and delayed (6-week) proteomic responses to DOX, commonly regulated proteins were identified via inner join on protein names. A bubble plot was constructed to visualize fold change relationships between timepoints, with points sized by combined magnitude of fold change and colored by directionality. GSEA was performed on both timepoints to identify significantly perturbed molecular pathways.

### 2.7. Analysis of Clinical Samples

To assess the translational relevance of our experimental findings, plasma samples were collected from breast cancer patients undergoing DOX-based chemotherapy. Samples were obtained at baseline and 24 hours after DOX infusion. These specimens were obtained prospectively as part of a randomized clinical study (NCT00895414) to examine the potential drug–drug interaction between DOX and enalapril, with institutional approval from both the Institutional Review Board and the Cancer Protocol Review Committee. Details of the primary study were described by Blaes et al. (11). Briefly, the study included breast cancer patients (N = 17 per arm) over the age of 18 with normal liver and kidney function, diagnosed with stage I–III breast cancer and receiving adjuvant doxorubicin (60 mg/m²) and cyclophosphamide (600 mg/m²) every 14 days for four cycles. In a crossover design, each patient received one cycle of chemotherapy with enalapril and another cycle without enalapril. For the present analysis, we exclusively utilized samples from participants in the non-enalapril treatment arm. Plasma levels of SERPINA3 were measured using commercially available Enzyme-Linked Immunosorbent Assay (ELISA) kit (catalog # EH411RB, Thermo Fisher Scientific) according to the manufacturer’s instructions.

### 2.8 Statistical Analysis

Statistical analyses for echocardiography, qPCR, western blotting, and ELISA data were performed using appropriate tests (Student’s t-test) with significance set at p < 0.05. Data are presented as mean ± standard error of the mean (SEM).

## 3. Results

### 3.1. Transcriptomic Alterations Induced by DOX in Mouse Cardiac Tissue

We previously demonstrated that chronic DOX exposure induced significant reductions in ejection fraction and fractional shortening, as assessed by echocardiography and originally presented in our published work (9). To understand the molecular mechanisms of DOX-induced cardiac dysfunction, bulk RNA-seq analysis was performed on cardiac tissue. This revealed significant alterations in the transcriptome following chronic DOX treatment (Figure 1). PCA demonstrated clear separation between the transcriptomic profiles of DOX-treated hearts and saline controls (Figure 1B). The first principal component (PC1) accounted for 53.32% of the total variation, indicating a robust and consistent treatment effect.

Differential gene expression analysis identified 64 significantly altered genes (27 upregulated and 37 downregulated) based on a threshold of |logLJ fold change| > 0.5 and adjusted p-value < 0.05, as visualized in a volcano plot (Figure 1C). Key upregulated genes included Nuclear Receptor Subfamily 1 Group D Member 1 (*Nr1d1*), Pleckstrin Homology Like Domain Family A Member 3 (*Phlda3*), Epoxide Hydrolase 1 (*Ephx1*), Transformation Related Protein 53 Inducible Nuclear Protein 1 (*Trp53inp1*), and D-box binding protein (*Dbp*). These genes are involved in processes like circadian rhythm/metabolism (*Nr1d1*), and apoptosis/stress response (*Phlda3, Trp53inp1*). Conversely, Apelin (*Apln*), a known cardioprotective peptide, and *Cd74* were identified as significantly downregulated, suggesting a loss of protective mechanisms and potential immune modulation.

A heatmap of the top 50 DEGs, arranged based on adjusted p-value, illustrated distinct expression patterns across individual samples, confirming consistent changes within treatment groups (Figure 1D). The upregulated genes clustered into functional categories including cardiac remodeling (Myosin Heavy Chain 7; *Myh7*), stress response (*Phlda3, Trp53inp1*), circadian regulation (*Nr1d1, Cirbp*), and xenobiotic metabolism (*Ephx1*). In addition, *Thbs2* (thrombospondin 2) upregulation suggests activation of fibrotic pathways, while increased expression of metabolic regulators *Hk1* (hexokinase 1), *Slc38a1*, and *Slc6a6* (amino acid transporters) indicates significant metabolic reprogramming. Among the downregulated genes, we observed coordinated suppression of immune response pathways, with multiple MHC class II genes (*H2-DMb1, H2-Aa, H2-Ab1, H2-Eb1*) and their master regulator *Ciita* showing reduced expression. This pattern, together with downregulation of *Cd74* and *Csf1r* (colony stimulating factor 1 receptor), indicates suppression of antigen presentation pathways in the heart following DOX treatment. Additionally, cardiac contractile function genes (*Ryr2, Xirp2*) and cardioprotective factors (*Apln, Tmsb4x*) were downregulated, suggesting compromised cardiac function and repair mechanisms. Among other downregulated genes were hemoglobin genes (*Hbb-bs, Hba-a1*), suggesting altered oxygen transport capacity; mitochondrial tRNA genes (*mt-Tc, mt-Tq*), whose reduced expression was consistent with DOX-induced mitochondrial toxicity; and *Mki67*, a marker of cell proliferation, indicating decreased cardiac regenerative capacity following DOX treatment.

GSEA identified significantly perturbed pathways in DOX-treated hearts (Figure 1E). Among the most significantly upregulated pathways were epithelial mesenchymal transition (EMT), myogenesis, P53 pathway, and cholesterol homeostasis, suggesting activation of stress responses, cellular transformation, and metabolic remodeling. The strong enrichment of the p53 pathway corroborates our observation of upregulated p53 target genes in the differential expression analysis. Conversely, pathways related to interferon gamma response, interferon alpha response, allograft rejection, and E2F Targets, were significantly downregulated, indicating suppression of immune responses and cell cycle progression.

### 3.2. qPCR Validation of Differentially Expressed Genes (DEGs) and Circadian Clock Dysregulation in DOX-treated Hearts

To validate and further investigate the transcriptomic findings, we performed qPCR analysis of key genes in cardiac tissue from DOX-treated and control mice. Validation of the RNA-seq findings using qPCR confirmed the significant upregulation of *Trp53inp1* (Figure 2A), *Ephx1* (Figure 2B), and *Slc6a6* (Figure 2C) mRNA levels, and the significant downregulation of *Cd74* (Figure 2D) and *Csf1r* (Figure 2E) mRNA in DOX-treated hearts compared to controls. This independent verification strengthens the transcriptomic results for several key genes.

**Figure 2.**
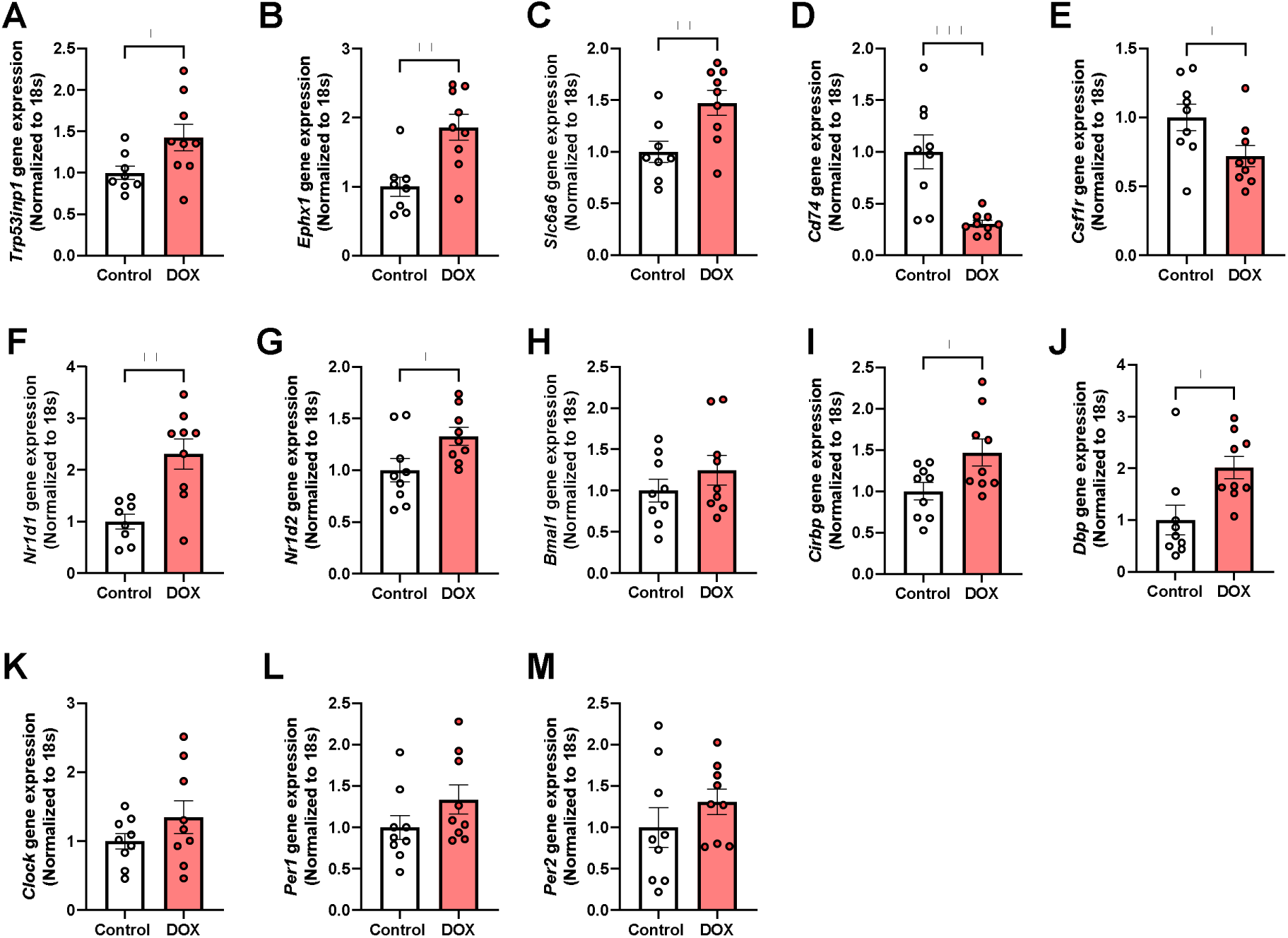
qPCR Validation of Differentially Expressed Genes (DEGs) and Circadian Clock Dysregulation in DOX-treated Hearts. (A-E) Validation of key DEGs identified by RNA-seq by qPCR: (A) *Trp53inp1*, (B) *Ephx1*, (C) *Slc6a6*, (D) *Cd74,* and (E) *Csf1r*. (F-M) Comprehensive analysis of circadian clock gene expression: (F) *Nr1d1* (Rev- erbα), (G) *Nr1d2* (Rev-erbβ), (H) *Bmal1*, (I) *Cirbp*, (J) *Dbp*, (K) *Clock*, (L) *Per1,* and (M) *Per2*. All gene expression values were normalized to 18S rRNA. Data are presented as mean ± SEM; n=9 per group; *p < 0.05, **p < 0.01, ***p < 0.001 vs. control group; unpaired t-test.

Furthermore, given the potential role of *Nr1d1* (also known as Rev-erbα, and identified in RNA-seq), a critical negative regulator of the circadian feedback loop, and the increasing recognition of circadian disruption in heart disease, we specifically examined the expression of key circadian clock genes. qPCR analysis revealed significant alterations in the expression profiles of several core clock components in the hearts of DOX-treated mice. The nuclear receptors *Nr1d1* (Rev-erbα, Figure 2F) and *Nr1d2* (Rev-erbβ, Figure 2G), which function as transcriptional repressors within the circadian feedback loop, were both significantly upregulated in DOX-treated hearts. These nuclear receptors are known to repress the transcription of *Bmal1* (*Arntl*), a core transcriptional activator in the circadian clock. Nevertheless, there was no significant change in *Bmal1* expression in DOX-treated hearts (Figure 2H).

Notably, two clock-controlled genes showed significant upregulation in DOX-treated hearts: *Cirbp* (Figure 2I), which modulates circadian gene expression post-transcriptionally and *Dbp* (Figure 2J), which is a direct target of the CLOCK-BMAL1 complex and an important mediator of clock output. No significant change was demonstrated in the expression of *Clock* (Figure 2K), which forms a heterodimer with Bmal1 to drive circadian gene transcription, or the Period genes *Per1* (Figure 2L) and *Per* 2 (Figure 2M), which encode negative regulators of the circadian feedback loop, in DOX-treated hearts. Collectively, these qPCR results confirm and substantially extend our transcriptomic findings, demonstrating that DOX treatment induces a coordinated dysregulation of the cardiac circadian clock network.

### 3.3. Proteomic Alterations Induced by DOX in Mouse Cardiac Tissue

Complementary proteomic analysis of cardiac tissue identified numerous proteins whose expression levels were significantly altered by chronic DOX treatment (Figure 3). Similar to the transcriptomics, PCA demonstrated a clear separation between the proteomic profiles of DOX- treated and control hearts (Figure 3A) suggesting a profound alteration in cardiac protein expression induced by DOX. The volcano plot highlighted significantly altered proteins (Figure 3B), with key upregulated examples including Myh7, Serpin Family A Member 3N (Serpina3n), Thrombospondin-1 (Thbs1), Ephx1, and Phospholamban (Pln). Downregulated proteins included Serpin Family A Member 1E (Serpina1e), and Haptoglobin (Hp).

**Figure 3.**
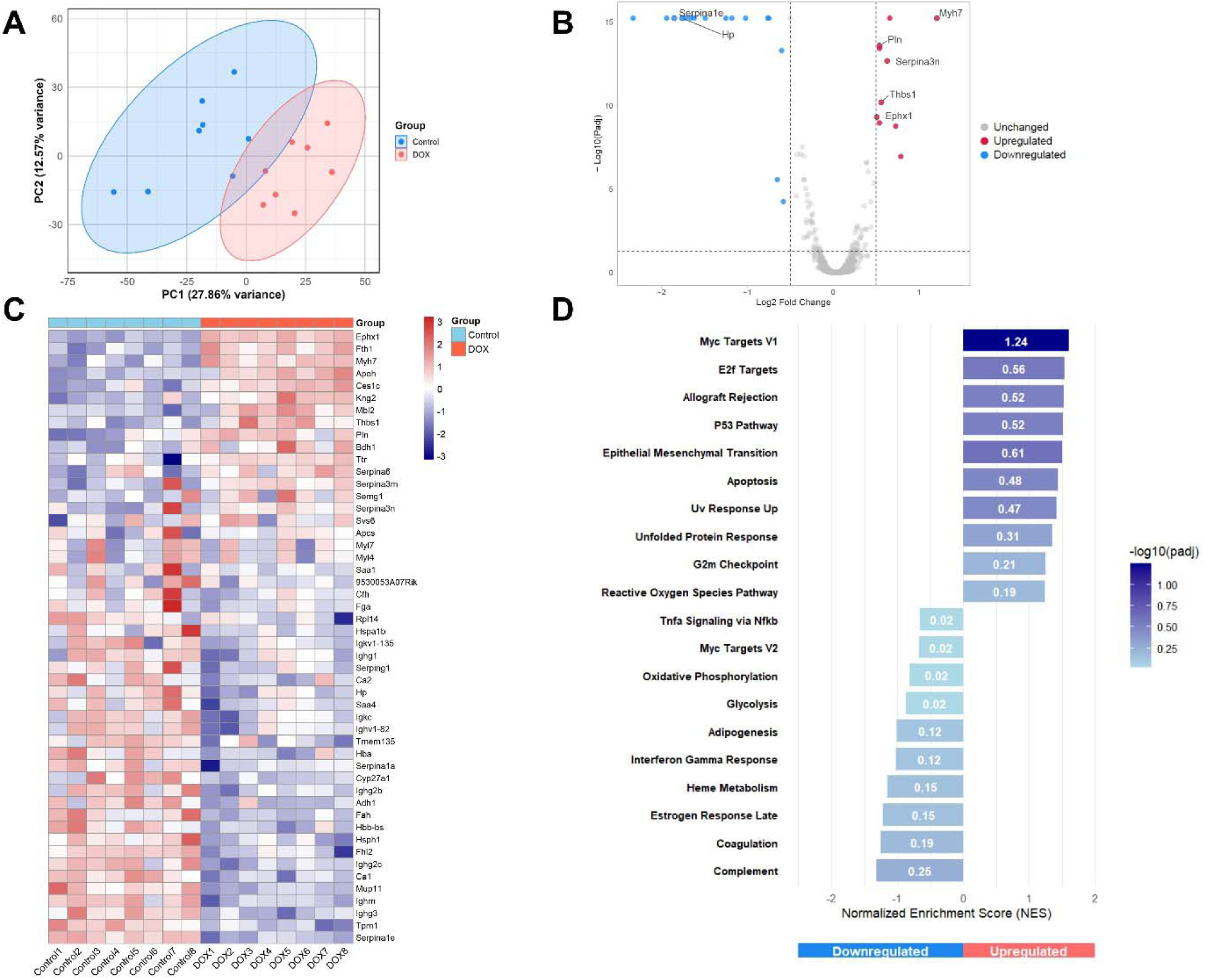
Proteomic Alterations Induced by DOX in Mouse Cardiac Tissue. (A) Principal Component Analysis (PCA) of cardiac proteome profiles from DOX-treated (red) and control (blue) mice. The ellipses represent 95% confidence intervals for each group. (B) Volcano plot showing differentially expressed proteins (DEPs) between DOX- treated and control hearts. The x-axis represents log2 fold change, and the y-axis represents -log10(p-value). Significantly upregulated proteins are shown in red, significantly downregulated proteins in blue, and proteins with non-significant changes in gray (based on |log_J fold change| > 0.5 and adjusted p-value < 0.05). Key DEPs are labeled. (C) Heatmap of the top 50 significantly DEPs in cardiac tissue from DOX-treated and control mice. Each column represents an individual sample, and each row represents a protein. Red indicates upregulation, blue indicates downregulation, and white indicates no change relative to the mean expression level. Two major clusters are evident: proteins upregulated in DOX-treated hearts (upper portion) and proteins downregulated in DOX-treated hearts (lower portion). (D) Gene Set Enrichment Analysis (GSEA) results showing selected significantly enriched pathways. Bars represent Normalized Enrichment Score (NES); positive NES indicates upregulation in DOX group, negative NES indicates downregulation. The color intensity of the bars corresponds to the -log10(padj) value, indicating the statistical significance of the enrichment.

Heatmap of the top 50 significantly DEPs, arranged based on adjusted p-value, revealed distinct expression patterns that clearly distinguished DOX-treated samples from controls (Figure 3C). Within the DOX-upregulated cluster, we observed enrichment of proteins involved in stress responses (Ephx1, Ces1c), iron sequestration (Fth1), acute phase responses (Apcs, Saa1), and protease inhibition (Serpina3n, Serpina3m, Serpina6). The upregulation of Pln suggests calcium handling dysregulation, while increased Bdh1 (3-Hydroxybutyrate Dehydrogenase 1) indicates a metabolic shift toward ketone utilization when mitochondrial function is compromised. The downregulated cluster contained proteins associated with cardiac contractile function (Tpm1, Fhl2), adaptive immunity (multiple immunoglobulins), mitochondrial function (Tmem135), and tissue protection (Serpina1a, Serpina1e, Hspb1). The downregulation of carbonic anhydrases (Ca1, Ca2) suggests altered pH regulation, while reduced Cyp27a1 indicates disrupted lipid metabolism. These patterns highlight coordinated DOX-induced proteomic remodeling in the heart.

GSEA identified significantly perturbed pathways in DOX-treated hearts (Figure 3D). Among the most significantly upregulated pathways were myc targets V1, E2F Targets, allograft rejection, P53 Pathway, and EMT, suggesting activation of stress responses, cell cycle regulation, and inflammatory processes. The strong enrichment of the p53 pathway is consistent with the known role of p53 in mediating DOX-induced cardiotoxicity and aligns with transcriptomics. Conversely, pathways related to complement, coagulation, estrogen response, heme metabolism, adipogenesis, oxidative phosphorylation, and interferon gamma response were significantly downregulated, indicating suppression of metabolic and immune processes. The downregulation of oxidative phosphorylation pathways is particularly noteworthy, as mitochondrial dysfunction is a well-established feature of DOX cardiotoxicity.

### 3.4. Elevation of Serpina3n/SERPINA3 in DOX-Treated Mouse Models and in Patients with Breast Cancer Receiving DOX

To assess the robustness of our proteomic findings and identify conserved biomarkers of DOX-induced cardiotoxicity, we compared our upregulated DEPs with those reported by Kruger et al. (12) who used the same dosing, but in a different mouse sub-strain (C57BL/6J). We observed a limited overlap in the proteomic response to DOX treatment (Figure 4A). The Venn diagram analysis revealed that two proteins, Myh7 and Serpina3n, were commonly upregulated in both studies, suggesting their potential relevance as conserved markers of cardiotoxicity. In addition, the limited overlap may reflect differences in genetic background between the C57BL/6N and C57BL/6J mouse sub-strains. Additionally, eight proteins were uniquely upregulated in our study while 11 proteins were uniquely upregulated in the Kruger et al. study based on criteria of |logLJ fold change| > 0.5 and adjusted p-value < 0.05. Given the consistent upregulation of Serpina3n across both mouse models, we further validated this finding by western blot analysis of cardiac tissue from our DOX-treated and control mice (Figure 4B). Consistent with our proteomic data, western blotting confirmed significant upregulation of Serpina3n protein levels in DOX-treated hearts compared to controls. To evaluate the potential translational relevance of our findings, we measured plasma concentrations of SERPINA3 (the human ortholog of mouse Serpina3n) in 17 breast cancer patients before and 24 hours after their DOX treatment (Figure 4C). Plasma SERPINA3 concentrations were significantly higher 24 hours post-DOX infusion compared to baseline. This rapid increase in circulating SERPINA3 following DOX administration suggests that SERPINA3 may serve as an early biomarker of DOX exposure in humans. Collectively, these findings highlight Serpina3n/SERPINA3 as a conserved response to DOX treatment across different mouse sub-strains and in human patients.

**Figure 4.**
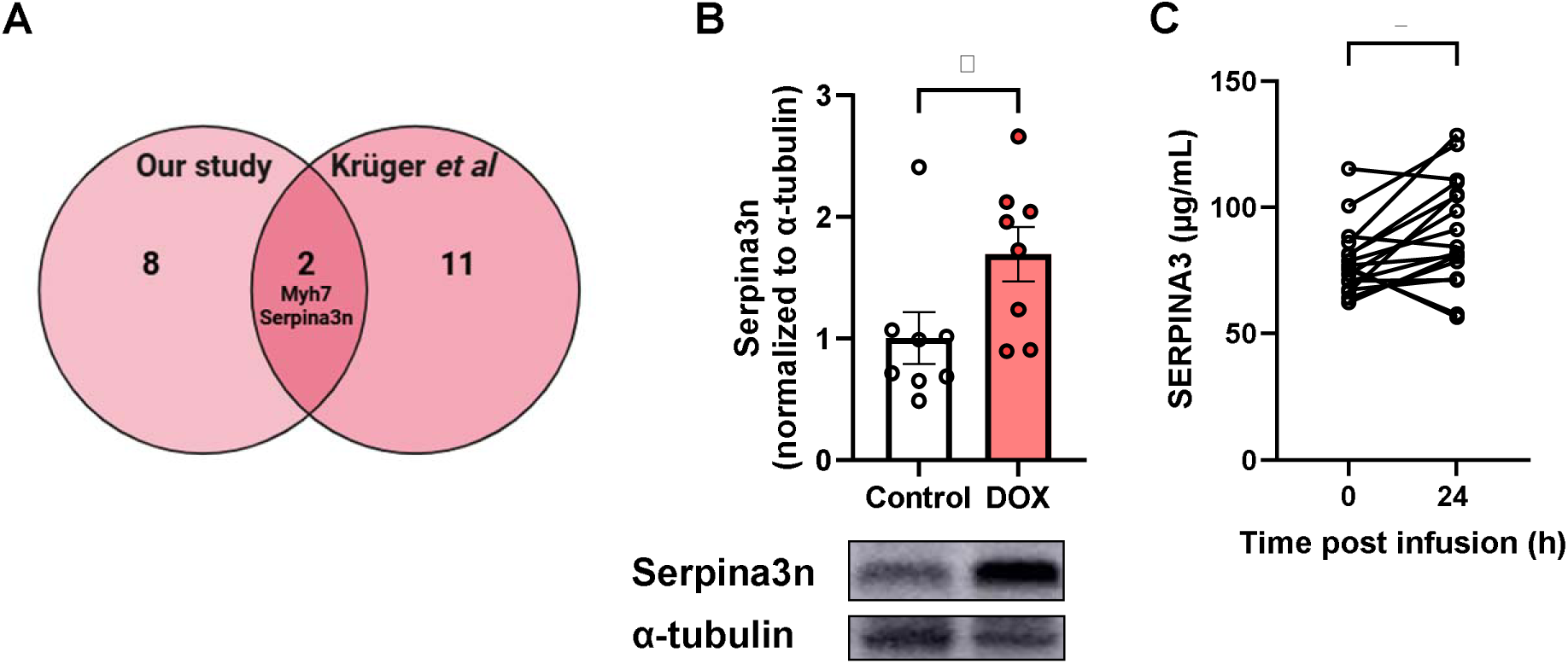
Serpina3n as a Biomarker of DOX Exposure Across Mouse Models and in Clinical Samples. (A) Venn diagram comparing upregulated differentially expressed proteins (DEPs) identified in our study with those reported by Kruger et al., highlighting Serpina3n and Myh7 as common upregulated DEPs. (B) Representative Western blot and quantification confirming significant upregulation of Serpina3n protein in mouse heart tissue from DOX-treated mice compared to controls (normalized to α-tubulin, mean ± SEM). (C) Plasma SERPINA3 (µg/mL) levels in breast cancer patients at baseline and 24 hours post-DOX infusion. Data are mean ± SEM. Individual data points represent biological replicates or patients. Asterisk denotes statistical significance (*p < 0.05).

### 3.5. Integrated Omics Analysis Between Transcriptomic and Proteomic Changes in DOX- Induced Cardiotoxicity

To gain a more comprehensive understanding of the molecular alterations underlying DOX-induced cardiotoxicity, we performed an integrated analysis of transcriptomic and proteomic data obtained from the same cardiac tissue samples (Figure 5). A Venn diagram illustrates the overlap between significantly altered genes (identified by RNA-seq) and proteins (identified by proteomics). Out of 13,067 entities identified by RNA-seq and 2,256 entities by proteomics, a total of 2,089 were found to be commonly altered at both the gene and protein levels, while 10,978 were unique to the transcriptome and 167 were unique to the proteome (Figure 5A).

**Figure 5.**
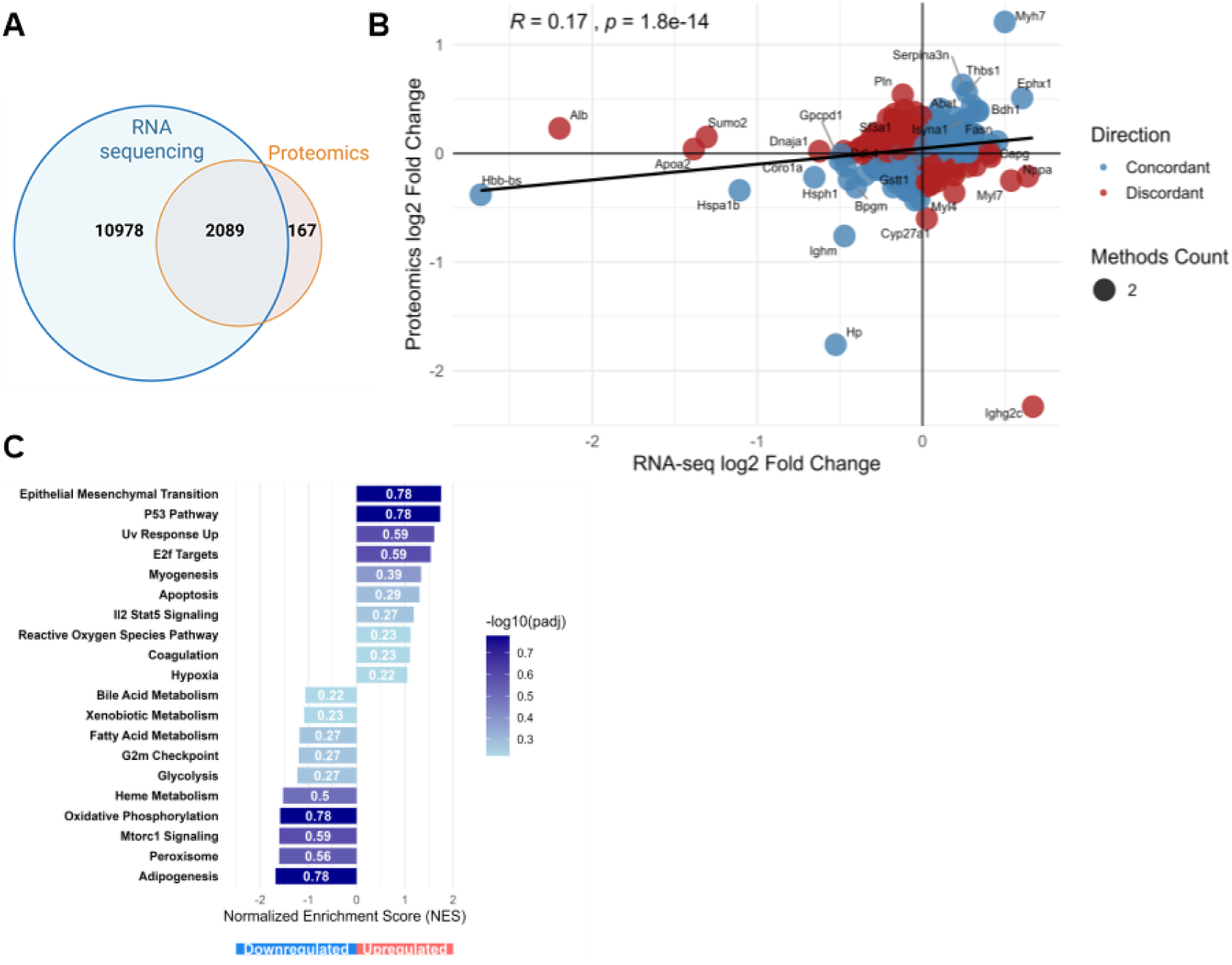
Integrated Omics Analysis Between Transcriptomic and Proteomic Changes in DOX-Induced Cardiotoxicity. (A) Venn diagram illustrating the overlap between altered transcripts (RNA-seq) and proteins (Proteomics). (B) Bubble plot correlating the log2 fold changes of commonly identified entities at the transcript (RNA- seq, x-axis) and protein (Proteomics, y-axis) levels. The correlation coefficient (R = 0.17) and p-value (p = 1.8e-14) are shown. Bubbles are colored based on concordance: blue for concordant changes (both mRNA and protein up- or downregulated) and red for discordant changes (mRNA and protein change in opposite directions). (C) Gene Set Enrichment Analysis (GSEA) performed on the concordantly regulated genes/proteins. The bar chart displays th Normalized Enrichment Score (NES) for significantly enriched pathways. Pathways enriched in downregulate concordant molecules are shown with negative NES values, and pathways enriched in upregulated concordant molecules are shown with positive NES values. The color intensity of the bars corresponds to the -log10(padj) value, indicating the statistical significance of the enrichment.

A bubble plot was generated for the commonly identified entities to visualize the correlation and concordance between mRNA (log2 Fold Change from RNA-seq) and protein (log2 Fold Change from proteomics) expression changes (Figure 5B). The plot shows a weak positive correlation between transcriptomic and proteomic changes (R = 0.17, p = 1.8e-14). This limited correlation, despite the highly significant p-value, suggests substantial post-transcriptional regulation in the cardiac response to DOX. Bubbles are colored to distinguish between concordant changes (both mRNA and protein up- or downregulated; N = 1060) and discordant changes (mRNA up/protein down, or vice-versa; N= 1029). Most prominently, Myh7 stands out as the most strongly upregulated molecule at both transcript and protein levels. Serpina3n, Thbs1, and Ephx1 displayed more pronounced upregulation at the protein level than at the transcript level, suggesting post-transcriptional enhancement. Conversely, Hbb-bs (hemoglobin beta), Hspa1b (heat shock protein family A member 1B), and Hp were downregulated at both levels. Notably, some molecules like Pln showed discordant regulation, being upregulated at the protein level but downregulated at the transcript level, while Ighg2c demonstrated the opposite pattern.

Pathway enrichment analysis was then performed specifically on the list of concordantly regulated genes/proteins (1060 entities) to identify biological processes robustly affected at both molecular levels (Figure 5C). Stress response pathways showed significant upregulation, with EMT and P53 pathway exhibiting the strongest enrichment, followed by UV response up and E2F Targets. Other upregulated pathways included myogenesis, apoptosis, IL2-STAT5 signaling, and reactive oxygen species Pathway, collectively indicating activation of stress responses, cell cycle regulation, and cell death mechanisms. Conversely, the analysis revealed significant downregulation of several metabolic pathways, including oxidative phosphorylation, adipogenesis, heme metabolism, glycolysis and fatty acid metabolism. The consistent downregulation of these energy metabolism pathways at both mRNA and protein levels underscores the profound impact of DOX on cardiac bioenergetics.

### 3.6. Temporal Comparison of Early and Delayed Molecular Responses to DOX-Induced Cardiotoxicity

To understand the temporal evolution of DOX-induced cardiotoxicity, we compared our early timepoint (4 days post-DOX) proteomics data with our previously published data by Abdelgawad et al.(10) representing a delayed timepoint (6 weeks post-DOX). The experimental timeline illustrates the administration of six IP DOX injections (4 mg/kg/week) over a period of six weeks, with tissue collection for proteomics analysis at two distinct timepoints: early (4 days after the last DOX dose) and delayed (6 weeks after the last DOX dose) (Figure 6A).

**Figure 6:**
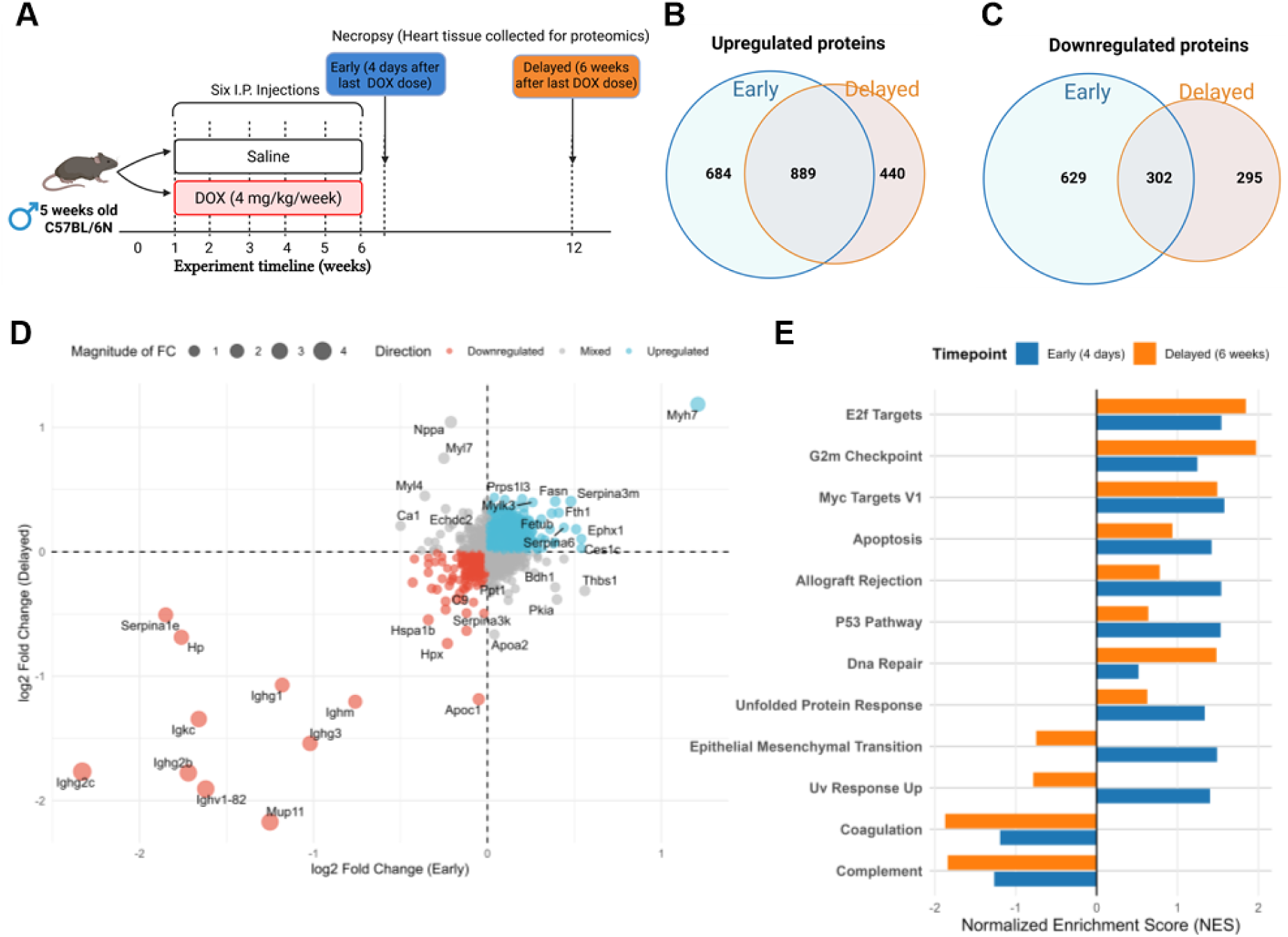
Temporal Comparison of Early and Delayed Molecular Responses to DOX-Induced Cardiotoxicity. (A) Experimental timeline showing the administration of six intraperitoneal DOX injections (4 mg/kg/week) to 5-week-old C57BL/6N mice over a period of six weeks. Heart tissue was collected for proteomic analysis at two distinct timepoints: early (4 days after the last DOX dose) and delayed (6 weeks after the last DOX dose). (B-C) Ven diagrams illustrating the overlap between differentially expressed proteins at early and delayed timepoints. (B) Upregulated proteins. (C) Downregulated proteins. (D) Scatter plot comparing the log2 fold changes of proteins at early (x-axis) and delayed (y-axis) timepoints. Proteins are colored according to their expression pattern: blue for upregulated, red for downregulated, and gray for mixed direction changes. The size of the bubbles represents th magnitude of fold change. Key proteins with notable temporal expression patterns are labeled. (E) Pathway enrichment analysis comparing early (blue bars) and delayed (orange bars) timepoints. The bar chart displays th Normalized Enrichment Score (NES) for significantly enriched pathways, with negative values indicatin downregulation and positive values indicating upregulation.

Venn diagram analysis of DEPs revealed substantial differences between the early and delayed responses to chronic DOX treatment. For upregulated proteins, we identified 684 proteins uniquely upregulated in the early timepoint and 440 proteins uniquely upregulated in the delayed timepoint, with 889 proteins consistently upregulated at both timepoints (Figure 6B). Similarly, for downregulated proteins, 629 proteins were uniquely downregulated in the early timepoint and 295 proteins uniquely downregulated in the delayed timepoint, with 302 proteins consistently downregulated across both timepoints (Figure 6C).

To further characterize the relationship between early and delayed protein expression changes, we generated a scatter bubble plot comparing the log2 fold changes at both timepoints (Figure 6D). This plot revealed several distinct patterns of temporal regulation. Proteins in the upper right quadrant (e.g., Myh7, Ephx1) showed sustained upregulation across both timepoints, suggesting their involvement in both early and delayed cardiotoxicity mechanisms. Proteins in the lower left quadrant (e.g., serpina1e, Hpx, Ighg2b, Ighv1-82) exhibited persistent downregulation, indicating sustained suppression of their associated functions. Notably, proteins in the upper left quadrant (e.g., Nppa) and lower right quadrant (e.g., Thbs1) demonstrated temporal switching in their expression patterns, being downregulated at one timepoint but upregulated at the other. This temporal switching suggests time-specific roles in the progression of cardiotoxicity.

Pathway enrichment analysis comparing the early and delayed timepoints revealed distinct temporal patterns in biological processes (Figure 6E). Several pathways showed stronger enrichment in the early timepoint, including unfolded protein response, P53 pathway, and allograft rejection, suggesting that acute stress responses, and inflammatory processes dominate the early response to DOX. This pattern suggests that acute stress responses are prominent immediately following DOX exposure but become attenuated over time, possibly due to resolution of acute injury or adaptation to chronic stress. Interestingly, the apoptosis pathway was enriched at both timepoints but more strongly in the early phase. In contrast, cell cycle and proliferation-related pathways such as E2F targets, G2M Checkpoint, and DNA Repair showed persistent but intensifying enrichment from early to delayed phases, indicating that cell cycle dysregulation and DNA damage responses become more prominent features of delayed chronic cardiotoxicity.

Notably, both complement and coagulation pathways exhibited consistent negative enrichment at both timepoints, with more pronounced downregulation at the delayed phase. This persistent suppression suggests that DOX induces sustained dysregulation of these homeostatic pathways, potentially contributing to chronic inflammatory and fibrotic responses throughout the progression of cardiotoxicity. Additionally, certain pathways showed opposing directions of enrichment between the two timepoints. EMT and UV Response Up pathways were upregulated in the early phase but downregulated in the delayed phase, suggesting that certain stress response mechanisms may be activated transiently and then suppressed as the heart adapts to chronic injury. This temporal comparison provides valuable insights into the dynamic nature of DOX-induced cardiotoxicity, revealing that different molecular mechanisms likely dominate at different stages of cardiac injury and remodeling.

## 4. Discussion

In this study, we employed an integrated-omics approach to comprehensively characterize the molecular landscape of DOX-induced cardiotoxicity in a mouse model and validated key findings in clinical samples. To our knowledge, this represents the first study to report integrated transcriptomics and proteomics analysis in response to DOX-induced cardiotoxicity in mice and to conduct temporal proteomics analysis comparing early and delayed molecular responses. Our integrated analysis of transcriptomic and proteomic data, coupled with temporal comparisons and clinical validation, has revealed several interconnected mechanisms underlying DOX cardiotoxicity.

### P53 Signaling and Stress Response

A prominent finding across our integrated-omics analyses was the consistent activation of p53 signaling pathways. Our transcriptomic data revealed significant upregulation of p53 target genes (*Phlda3, Trp53inp1*) and strong enrichment of the p53 pathway in GSEA analysis. This was corroborated at the protein level, where the p53 pathway was among the most significantly enriched in DOX-treated hearts. These findings align with mounting evidence that p53 serves as a master regulator of DOX-induced cardiotoxicity (13, 14). Recent transcriptomic profiling by McSweeney et al. similarly identified p53 as a key upstream regulator of DOX-induced transcriptional changes in cardiomyocytes (14).

The p53-dependent mechanisms of cardiotoxicity include induction of apoptosis, cell cycle arrest, and metabolic dysregulation (15). Our temporal analysis further revealed that p53 pathway activation is most pronounced during the early timepoint (4 days post-DOX), suggesting it has a primary role in mediating early cardiotoxic effects following DOX. This temporal pattern is consistent with recent findings demonstrating that early p53 activation represents a critical molecular event in the initial cardiac response to anthracycline exposure (13, 16). The identification of specific p53 effectors in our study, such as *Phlda3* (a p53- regulated repressor of Akt signaling) and *Trp53inp1* (a regulator of stress-induced autophagy), provides potential targets for intervention. Recent work showed that Phlda3 inhibits AKT survival signaling, amplifying cardiomyocyte loss (17, 18), and Trp53inp1 promotes autophagy-dependent cell death (19).

The role of p53 in DOX- induced cardiotoxicity appears to be temporally nuanced. While our findings corroborate earlier work demonstrating that p53 deletion or inhibition attenuates DOX-induced cardiac dysfunction and apoptosis in mice (20, 21), other studies have revealed a protective function of p53. In particular, p53 appears essential for maintaining long-term mitochondrial integrity and cardiac function, as its chronic deletion or inhibition has been shown to exacerbate mitochondrial dysfunction and increase cell death (22, 23). This dual role may explain why we observed the most pronounced p53 pathway activation during the early timepoint of cardiotoxicity.

### Circadian Rhythm Disruption

One of the intriguing findings in our study was the significant dysregulation of circadian clock genes, particularly the upregulation of repressors (*Nr1d1* and *Nr1d2*) observed in both transcriptomic and qPCR analyses and altered clock outputs (*Cirbp, Dbp*). The coordinated dysregulation of multiple clock components, including positive regulators, repressors, and output genes indicates a systematic disruption of the cardiac circadian network rather than isolated changes in individual genes. The circadian system plays a crucial role in maintaining cardiovascular homeostasis by regulating heart rate, blood pressure, and metabolic processes. It orchestrates daily rhythms in cardiac metabolism, contractility, and cellular repair, and its disturbance can exacerbate cardiovascular disease risk and impair recovery from injury (24, 25). Recent studies have shown that DOX persistently alters the expression and acetylation patterns of core circadian genes in the mouse heart, even weeks after drug exposure (26, 27). Consistent with our findings, a recent transcriptomic analysis in rats revealed activation of circadian rhythm signaling in response to DOX, including enrichment of *Nr1d1*, *Dbp*, and *Per* genes which may contribute to the worsening of lipid metabolism impairment (28). The upregulation of *Nr1d1* and *Nr1d2*, which function as transcriptional repressors within the circadian feedback loop, may have downstream effects on metabolic pathways and inflammatory responses. These nuclear receptors regulate genes involved in lipid metabolism, mitochondrial function, and inflammatory signaling (29), all of which are perturbed in DOX cardiotoxicity. Recent work demonstrated that pharmacological activation of Nr1d1 attenuates DOX-induced cardiotoxicity, suggesting a potential therapeutic approach (30).

### Metabolic Reprogramming and Mitochondrial Dysfunction

A consistent finding across our transcriptomic and proteomic analyses was the profound metabolic reprogramming in DOX-treated hearts. Our integrated omics analysis revealed concordant downregulation of multiple metabolic pathways, including oxidative phosphorylation, fatty acid metabolism, and heme metabolism. This systematic suppression of energy metabolism pathways underscores the central role of bioenergetic failure in DOX cardiotoxicity (31). DOX preferentially accumulates in cardiac mitochondria, promoting excessive ROS generation, which damages mitochondrial DNA and impairs electron transport chain activity (31). In our study, this is further evidenced by the downregulation of mitochondrial tRNA genes (*mt-Tc, mt-Tq*) and suppression of key metabolic enzymes and transporters. In response, the heart shifts toward glycolysis, as indicated by upregulation of *Hk1*, and increases amino acid transporter expression (*Slc38a1*, *Slc6a6*), reflecting an adaptive, yet ultimately insufficient, attempt to compensate for impaired oxidative metabolism. Proteomic data also show increased Bdh1, suggesting a shift toward ketone utilization, a phenomenon also observed in heart failure and recently linked to improved cardiac energetics in DOX models (32, 33). Additionally, downregulation of carbonic anhydrases (*Ca1, Ca2*) and *Cyp27a1* point to disrupted pH and lipid homeostasis.

These findings align with recent metabolomic studies demonstrating that DOX induces profound metabolic perturbations in cardiac tissue, characterized by impaired fatty acid oxidation (34, 35). Collectively, these findings underscore that DOX-induced metabolic reprogramming and mitochondrial dysfunction are tightly interconnected, driving a cycle of energy depletion, oxidative stress, and progressive cardiac dysfunction. Therapeutic strategies aimed at restoring mitochondrial function, such as iron chelation with dexrazoxane, antioxidant therapy (e.g., melatonin, MitoQ), mitophagy enhancement (metformin), and metabolic support with ketones or SGLT2 inhibitors have shown promise in experimental models and represent potential avenues for mitigating DOX cardiotoxicity (33, 36, 37).

### Protease Inhibitor Dysregulation and Serpina3n as a Biomarker

An interesting finding in our proteomic analysis was the significant upregulation of multiple serpin family members, particularly Serpina3n, and downregulation of serpina1e in DOX-treated hearts. Our comparative analysis with proteomic data from Kruger et al. (12) revealed Serpina3n as one of only two proteins consistently upregulated across different mouse strains, suggesting it represents a conserved response to DOX. Most importantly, the higher plasma SERPINA3 concentrations 24 hours post-DOX infusion suggests it should be evaluated as an early biomarker of DOX exposure and possible predictor of cardiotoxicity risk.

SerpinA3n, the murine homologue of human SERPINA3 (α-1 antichymotrypsin), has emerged as a potential biomarker and potential mechanistic contributor in DOX-induced cardiotoxicity. This acute-phase reactant and serine protease inhibitor is consistently upregulated in cardiac and vascular tissue and plasma following DOX exposure, with expression predominantly localized to endothelial cells and cardiomyocytes (38). Notably, SerpinA3n shows sustained elevation in cancer survivors experiencing cardiovascular toxicity after DOX therapy cessation and demonstrates an inverse correlation with left ventricular ejection fraction (12). Beyond its utility in DOX-induced cardiotoxicity, SerpinA3n has been suggested as a marker for heart failure and adverse outcomes in acute myocardial infarction, with increased levels associated with poor clinical prognosis (39, 40). Its functional significance has been partially elucidated in a recent study on myocardial infarction, where genetic deletion exacerbates post-infarct cardiac dysfunction and inflammation, suggesting a cardioprotective role (41). However, despite the accumulating evidence from multi-omics studies and translational research establishing SerpinA3n/SERPINA3 as a robust biomarker of cardiac injury, its precise mechanistic role in DOX-induced cardiotoxicity remains incompletely understood. Future investigations should focus on delineating the molecular pathways through which SerpinA3n influences cardiac responses to DOX exposure, potentially revealing novel therapeutic targets for preventing or mitigating this serious complication of cancer therapy.

### Cardiac Remodeling and Contractile Dysfunction

Our multi-omics approach revealed molecular mechanisms underlying the structural and functional deterioration observed in DOX-induced cardiac dysfunction. Chronic DOX exposure induced significant systolic dysfunction (reduced EF, FS, and CO) and structural remodeling, including ventricular dilation and wall thinning. These functional abnormalities correspond with transcriptomic and proteomic evidence of impaired contractility and disrupted calcium homeostasis. Specifically, DOX downregulated genes critical for cardiac contractile function, such as *Ryr2, Xirp2*, and *Tpm1*, while upregulating *Myh7*, indicating a reactivation of the fetal gene program, which is a hallmark of pathological remodeling. In addition, DOX upregulated Pln, suggesting calcium handling abnormalities that may contribute to the observed systolic dysfunction. Pln is a regulatory protein that modulates the activity of sarcoplasmic reticulum Ca²⁺-ATPase 2a (SERCA2a), where dephosphorylated Pln inhibits the affinity of the SERCA2a for calcium and phosphorylation relieves this inhibition (42).

The downregulation of Ryr2 likely contributes to impaired excitation contraction coupling, consistent with findings that DOX-and daunorubicin-induced Ryr2 dysfunction leads to calcium handling abnormalities (43, 44). Notably, we observed downregulation of cardioprotective factors *Apln* and *Tmsb4x* (thymosin beta-4), suggesting compromised endogenous repair mechanisms. *Apln* has established cardioprotective properties, and its downregulation may exacerbate DOX cardiotoxicity (45, 46). Tmsb4x/Tβ4 is a potent endogenous cardioprotective factor that mitigates oxidative stress, inflammation, and apoptosis (47, 48). Although its direct role in DOX-induced cardiotoxicity remains to be fully elucidated, substantial mechanistic overlap strongly suggests that reduced Tmsb4x expression may worsen cardiac injury, and therapeutic augmentation of Tβ4 could be beneficial (49).

### Thrombospondins and Fibrotic Remodeling

Upregulation of *Thbs2* (thrombospondin-2) in transcriptomics, Thbs1 in proteomics, and enrichment of Epithelial-Mesenchymal Transition pathway in transcriptomic, proteomic, and concordant integrated omics imply fibroblast activation, endothelial dysfunction, and extracellular matrix (ECM) deposition, consistent with early fibrotic remodeling. Thbs1 and Thbs2 are matricellular glycoproteins that modulate ECM composition, angiogenesis, and cellular responses to stress (50). In DOX-induced cardiotoxicity, their upregulation may reflect a compensatory response to myocardial injury. Notably, the absence of Thbs2 has been associated with increased cardiomyocyte damage and matrix disruption in DOX-induced cardiomyopathy, indicating a protective role for Thbs2 in maintaining ECM integrity (51).

The enrichment of the Epithelial-Mesenchymal Transition pathway implies that DOX exposure may induce phenotypic changes in cardiac cells, particularly endothelial cells transitioning to mesenchymal-like states, a process known as endothelial-to-mesenchymal transition (EndMT). This transition contributes to fibrosis and impaired cardiac function (52). Studies have highlighted the involvement of EndMT in DOX-induced cardiotoxicity, suggesting that targeting this pathway could mitigate adverse cardiac remodeling (53). Furthermore, interventions targeting EndMT, such as calcitriol, have been shown to attenuate DOX-induced cardiac fibrosis and improve cardiac function by inhibiting the EndMT process (54).

### Immune Modulation

The coordinated downregulation of multiple immune-related genes, including MHC class II molecules (*H2-DMb1, H2-Aa, H2-Ab1, H2-Eb1*), their master regulator *Ciita*, and *Cd74* indicates suppression of antigen presentation and macrophage-dependent repair in the heart, potentially impairing the clearance of damaged cells and limiting tissue repair. This immune paralysis, coupled with reduced proliferative capacity (*Mki67* downregulation), likely hinders clearance of damaged cells and tissue regeneration. At the protein level, we observed differential regulation of immune components, with upregulation of acute phase proteins (Saa1, Apcs) alongside downregulation of multiple immunoglobulin components (Ighg1, Ighg2b, Ighg2c, Ighm, Ighg3). Our pathway analysis further supported immune modulation as a key feature of DOX cardiotoxicity, with downregulation of interferon response pathways. These findings are consistent with recent studies demonstrating that DOX induces significant immune abnormalities and inflammatory responses that contribute to progressive cardiovascular damage (55, 56). The downregulation of *Cd74*, which also functions as a receptor for macrophage migration inhibitory factor (MIF), may compromise MIF-mediated cardioprotection, as suggested by Xu et al. (57). Our findings regarding *Cd74* are consistent with a recent study (58), which demonstrated that DOX treatment leads to significant downregulation of Cd74 in both male and female mice, with a more pronounced effect observed in males.

In conclusion, our comprehensive integrated multi-omics approach has revealed the complex molecular landscape underlying DOX-induced cardiotoxicity, characterized by p53 pathway activation, circadian rhythm disruption, metabolic reprogramming, contractile dysfunction, and mitochondrial dysfunction. The identification of Serpina3n/SERPINA3 as a conserved DEP across mouse models and in human patients provides compelling rationale for further investigation of its potential as an early detection biomarker for cardiotoxicity. The temporal dynamics of these molecular alterations, with distinct signatures in early versus delayed phases, suggests that therapeutic interventions may need to be tailored to specific timepoints in the progression of cardiac injury. Future studies should focus on further evaluating these potential biomarkers in larger clinical cohorts to establish their predictive utility and exploring targeted interventions against key pathways, particularly p53 signaling, circadian disruption, and metabolic dysfunction, to develop effective cardioprotective strategies that preserve the therapeutic efficacy of DOX while minimizing its detrimental cardiac effects.

### Clinical Perspectives

- **Background:** Doxorubicin is a widely used chemotherapeutic agent whose clinical utility is significantly constrained by cardiotoxicity. The precise molecular mechanisms underlying doxorubicin-induced cardiotoxicity remain incompletely understood, hindering the development of effective cardioprotective strategies.
- **Brief summary of the results:** Our integrated multi-omics approach revealed key mechanisms of doxorubicin cardiotoxicity, including p53 pathway activation, circadian rhythm disruption, metabolic reprogramming, mitochondrial dysfunction, and immune modulation, with SERPINA3 as a potential biomarker in breast cancer patients receiving DOX.
- **Potential significance of the results to human health and disease:** Identification of early molecular signatures and temporal dynamics of DOX-induced cardiotoxicity provides opportunities for biomarker development and targeted interventions, that could prevent cardiac damage while preserving anticancer efficacy.

## Data Availability

Data will be made available on request.

## CRediT Author Contribution

**Mohamed Dabour:** Methodology, Conceptualization, Data curation and analysis, Writing— original draft**. Ibrahim Abdelgawad, Bushra Sadaf, Mary R. Daniel, Marianne Grant:** Methodology, Conceptualization, Writing – review & editing**. Anne Blaes, Pamala Jacobson:** Investigation, Resources, Writing – review & editing. **Beshay Zordoky:** Conceptualization, Funding acquisition and supervision, Writing – review & editing.

## Ethics Approval

All animal procedures were approved by the Institutional Animal Care and Use Committee at the University of Minnesota (Protocol ID: 2106–39176 A) and conducted in accordance with relevant guidelines.

## Supporting information

RNA sequencing data

Proteomics data

## Acknowledgments

We thank LeeAnn Higgins, Todd Markowski, and Cesar Anguaya Velasquez in the Center for Metabolomics and Proteomics at the University of Minnesota for providing services related to the generation of quantitative proteomics data. The Orbitrap Eclipse instrumentation platform used in this work was purchased through High-end Instrumentation Grant S10OD028717 from the NIH. Experiments using the Amersham Imager were supported by the resources and staff at the University of Minnesota University Imaging Centers (UIC).

## Abbreviations

DOX: Doxorubicin
DEGs: Differentially Expressed Genes
DEPs: Differentially Expressed Proteins
RNA-seq: RNA sequencing
TMT: Tandem Mass Tag
GSEA: Gene Set Enrichment Analysis
PCA: Principal Component Analysis
ECM: Extracellular Matrix
ROS: Reactive Oxygen Species
qPCR: Quantitative Polymerase Chain Reaction
FDR: False Discovery Rate
SERPINA3: Serpin Family A Member 3

